# A reversible feedback mechanism regulating mitochondrial heme synthesis

**DOI:** 10.1101/2025.06.09.658730

**Authors:** Iva Chitrakar, Alexis B. Roberson, Pedro H. Ayres-Galhardo, Breann L. Brown

## Abstract

Proper heme biosynthesis is essential for numerous cellular functions across nearly all life forms. In humans, dysregulated heme metabolism is linked to multiple blood diseases, neurodegeneration, cardiovascular disease, and metabolic disorders. Erythroid heme production begins with the rate-limiting enzyme Aminolevulinic Acid Synthase (ALAS2) in the mitochondrion. Although prior studies discuss the regulation of ALAS2 in the nucleus and cytoplasm, its modulation as a mature mitochondrial matrix enzyme remains poorly understood. We report that heme binds mature human ALAS2 with high affinity, acting as a reversible mixed inhibitor that reduces enzymatic activity. Structure-based modeling reveals two flexible regions of ALAS2 interact with heme, locking the enzyme in an inactive conformation and occluding the active site. Our work reveals a negative feedback mechanism for heme synthesis, offering insights into the spatial regulation of ALAS2 and the maturation of the essential heme cofactor.

## Introduction

Our knowledge of cellular metabolism continues to expand with our understanding of the role of metabolites as cell signaling effectors (reviewed in ^1^). Consequently, metabolic processes, which form the foundation of organismal biology, are studied in a new context to reveal critical insights into chemical and cellular signaling. Heme metabolism is central to multiple biological processes as it is a vital cofactor supporting oxygen transport, transcriptional regulation, and drug detoxification, among others^2-4^. Heme production must be tightly controlled due to the toxicity associated with heme deficiency or excessive amounts of porphyrin precursors, which produce reactive oxygen species and free radicals^5-8^.

The enzyme that catalyzes the rate-limiting step for heme biosynthesis in many organisms is aminolevulinic acid synthase (ALAS)^9,10^, a PLP-dependent homodimeric enzyme that mediates the condensation of glycine and succinyl-CoA to produce aminolevulinic acid^11,12^. Humans contain two ALAS isoforms, ALAS1 and ALAS2^13-15^. ALAS1 is a ubiquitously expressed housekeeping enzyme ^13,14^, and ALAS2 is the erythroid-specific isoform that governs the synthesis of 85-90% of the total body heme pool^5,16^. The high demand for heme to produce hemoglobin implies that erythroid heme synthesis is not tightly regulated. However, the fact that multiple blood disorders arise from dysfunctional heme biosynthesis indicates otherwise. Current clinical reports have identified over 95 human *ALAS2* mutations that result in the loss-of-function disease X-linked sideroblastic anemia (XLSA)^17^ or the gain-of-function disease X-linked protoporphyria (XLPP)^18,19^. Compelling new evidence reveals that *ALAS2* is also expressed in non-erythroid cell^20^, underscoring our critical need to understand ALAS2 enzyme regulation for metabolic processes beyond erythropoiesis.

Previous research primarily focused on the nuclear and cytosolic regulation of ALAS^21-31^. For example, heme binds directly to the ALAS mitochondrial targeting sequence, preventing protein translocation into the matri^21,22^. However, little is known about its enzymatic modulation in the mitochondrial matrix, which is the cellular compartment where heme biosynthesis is initiated and terminated. Human ALAS paralogs contain multiple putative heme-binding motifs in the mature enzyme, which lacks the mitochondrial targeting sequence. Various reports established that the interaction between heme and mitochondrial ALAS1 results in either LONP1 or CLPXP-mediated degradation^25,32,33^. Further, ALAS homologs from various α-proteobacteria are reported to be inactivated by heme binding^34,35^. Whether human ALAS2 is subject to negative regulation by heme in mitochondria remains to be fully explored.

Due to the complex impacts of heme variation on effective erythropoiesis, gene transcription, and cseveral other cell signaling pathways, it is essential to establish *in vitro* principles of enzyme regulation that form the foundation of interpreting physiological outcomes.

Here, we identified a reversible mechanism by which heme inhibits its synthesis by affecting mature mitochondrial ALAS2 activity, challenging the notion that erythroid heme synthesis is an all-or-nothing process. This mechanism provides a response to heme stress that rapidly inactivates ALAS2, thereby decreasing heme synthesis. With our updated understanding of ALAS2 tissue expression, our study details a new mechanism of negative feedback regulation for heme biosynthesis, regulating erythropoiesis and potentially the maturation of hemoproteins, controlling numerous biological functions.

## Results

### The human ALAS2 sequence contains multiple heme-binding motifs

One challenge to identifying transient or regulatory heme-protein interactions is the diversity of interaction motifs and amino acid ligands. Although many heme-binding proteins use histidine residues as the axial ligand to coordinate the heme iron, other examples employ cysteine, methionine, tyrosine, or lysine residues^36^. The lower abundance of non-histidine motifs makes predicting these sites less accurate than other canonical motifs^37-39^. However, certain parameters have been identified to aid in prediction.

Previous studies identified a heme regulatory motif (HRM) that consists of a core cysteine-proline (CP) di-peptide in human ALAS1, with cysteine functioning as the axial ligand to the heme iron^21^. A multiple sequence alignment of vertebrate ALAS enzymes revealed the presence of five CP motifs conserved in both isoforms across homologs (**Fig. 1a, Supplemental Fig. 1**). The first two N-terminal HRMs reside in the mitochondrial target sequence, and heme binding to these sites negatively regulates ALAS mitochondrial import^21,22^. The remaining three motifs are dispersed throughout the N-terminal extension and catalytic core of ALAS (**Fig. 1b**).

**Figure 1.**
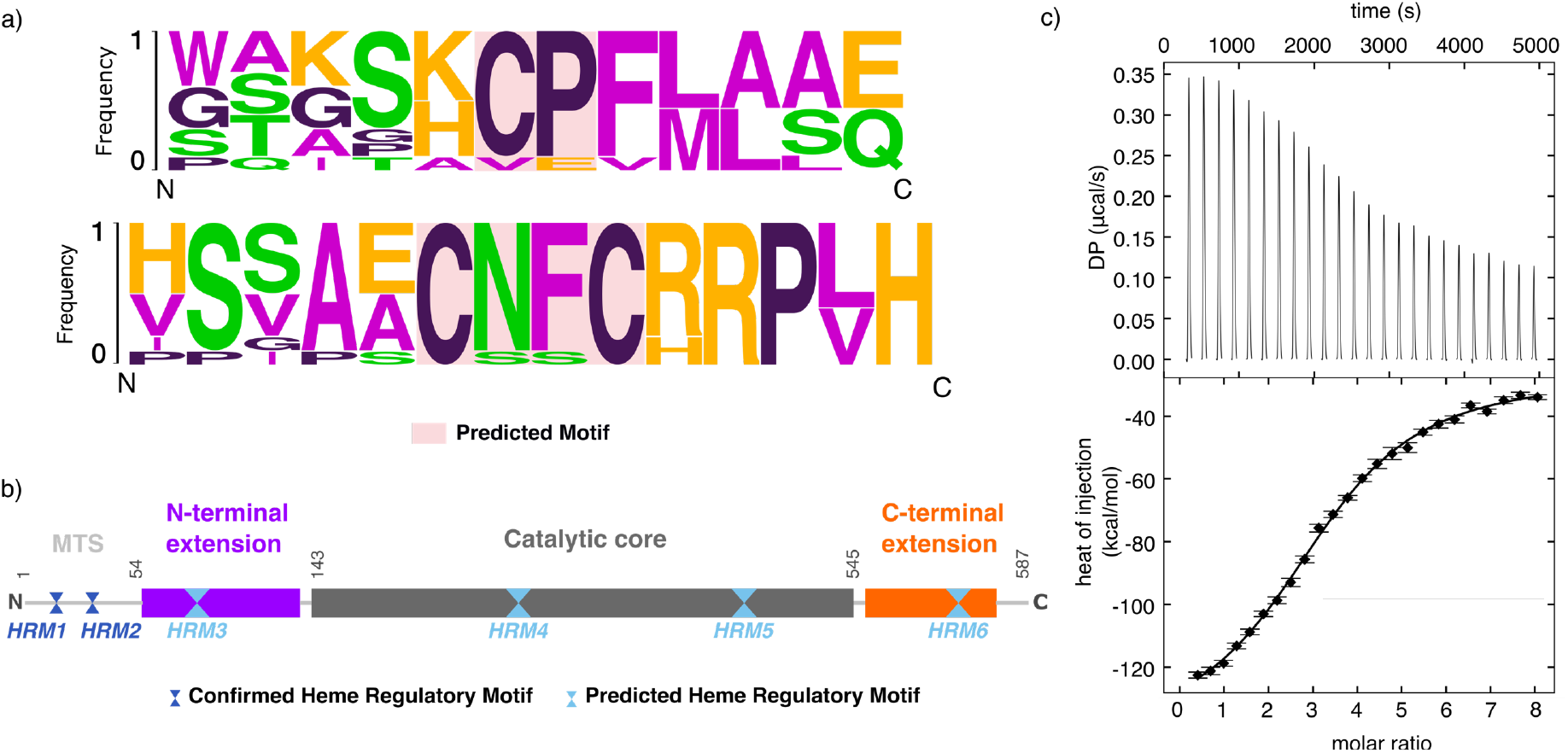
Conserved heme regulatory motif in ALAS. (a) Consensus sequence logo generated based on the heme regulatory motifs (HRM) conserved across multiple ALAS homologs. Conserved motifs (pink box) include CP dipeptide and CXXC. (b) Domain map of human ALAS2 highlighting the experimentally confirmed HRMs in the mitochondrial target sequence (MTS) and predicted HRMs in dark and light blue, respectively. The ALAS N-terminal extension is colored purple and the C-terminal extension is colored orange. The motifs according to human ALAS2 residue numbers are HRM1: ^11^CP^12^; HRM2: ^38^CP^39^; HRM3: ^70^CP^71^; HRM4: ^338^CP^339^; HRM5: ^471^CP^472^; and HRM6: ^555^CXXC^558^. (c) Representative ITC thermograph (top) and binding isotherm (bottom) for the binding between hemin and WT human ALAS2.

In addition to the CP motif, a variation of a separate conserved heme-binding motif commonly found in cytochromes was also identified. This canonical CXXCH sequence, where X represents any amino acid, is found in the human ALAS2 C-terminal domain as CXXCR. Interestingly, the mouse homolog contains the classical CXXCH motif in the same position. In contrast, many other vertebrate ALAS enzymes retain the Arg residue in the terminal position (**Fig. 1a**). Collectively, ALAS2 contains two verified and four predicted HRMs per protomer, referred to as HRM1 through HRM6 in this paper.

### *Human ALAS2 binds heme* in vitro

Due to the various motifs present in human ALAS2, we sought to determine if there is a role for heme in regulating mature ALAS2 after it is processed in the mitochondrial matrix. Mature wildtype ALAS2 (WT, residues 54-587), which lacks the mitochondrial targeting sequence, was expressed and purified to homogeneity. Titration of WT ALAS2 into hemin (ferric chloride heme) resulted in robust binding exhibited by an exothermic binding curve (**Fig. 1c**). The ALAS2-heme dissociation constant was determined to be ∼230 nM (**Table 1**). The binding stoichiometry was best fit to 4:1 (monomer equivalents), corresponding to two heme binding sites per ALAS2 protomer. Thus, heme binds mature ALAS2 tightly, and potentially at multiple sites.

**Table 1.** Isothermal Titration Calorimetry thermodynamic parameters.

| Construct | K<sub>d</sub> (nM) | ΔH (kcal/mol) | ΔS (kcal/mol*K) | ΔG (kcal/mol) |
| --- | --- | --- | --- | --- |
| WT | 232 [170, 324] | -382 [-440, -342] | -1.25 | -9.05 |
| ΔN | 589 [322, 1428] | -1006 [-1432, -810] | -3.35 | -8.50 |
| ΔC | 279 [220, 360] | -544 [-604, -499] | -1.80 | -8.94 |
| Cys-mut | 610 [401, 1132] | -361 [-561, -280] | -1.18 | -8.48 |

\*95% confidence interval in brackets

### Heme binds reversibly to inhibit ALAS2 activity

Having established that there is robust heme binding to mature ALAS2, we sought to determine the functional impact of this interaction. The *in vitro* enzyme activity of ALAS2 was measured by monitoring the rate of ALA product release. WT ALAS2 activity decreased significantly with increasing hemin concentration, yielding an IC_50_ value of 18.7 ± 0.5 µM (**Fig. 2a, Table 2**). These studies were performed using the catalytically active holoenzyme that is covalently bound to the PLP cofactor. This covalent adduct is broken and reformed throughout the ALAS reaction cycle (**Supplemental Fig. 2**). To address the mechanism of heme inhibition, we generated a catalytically inactive form of ALAS2 by chemically cleaving the covalent bond between the endogenous PLP cofactor and a conserved active site Lys (apoALAS2). The apoenzyme is then reactivated with the addition of fresh cofactor. In the absence of heme, increasing concentrations of PLP restored enzymatic activity, with an activation constant (K_a_) of c0.72 ± 0.08 μM (**Fig. 2b**). In the presence of 50 μM heme, PLP partially rescued activity with a 7 ± 1 μM EC_50_ (**Fig. 2b**). It is possible that there is not complete restoration of maximal enzyme activity if a portion of apoALAS2 molecules contain damaged cofactor that is unable to be exchanged for exogenous PLP. These data reveal that heme-mediated inhibition is reversible. ALAS2 activity was monitored with increasing amounts of PLP in the presence of various heme concentrations, up to 100-fold excess. The resulting data revealed significant differences in maximal enzyme velocity (V_max_) and an increased PLP K_a_, indicating that heme binds as a mixed inhibitor that binds both the free and cofactor-bound enzyme (**Fig. 2c**). We also measured the activity of purified ALAS2 holoenzyme in response to variable heme and succinyl-CoA substrate concentrations (**Fig. 2d**). Increasing amounts of succinyl-CoA still rescue heme-dependent inhibition. However, heme binds uncompetitively to the ALAS2:succinyl-CoA holoenzyme complex rather than the free enzyme, leading to decreases in both enzyme velocity (8-fold lower V_max_) and substrate affinity (3-fold lower EC_50_,**Fig. 2e**). Due to the sequential nature of the ALAS2 reaction, we propose that the uncompetitive inhibition results from diminished free enzyme available at this stage of the catalytic cycle. Combined, these data establish that heme acts as an inhibitor of mature human ALAS2, which can bind allosteric sites to decrease enzyme activity.

**Table 2.**
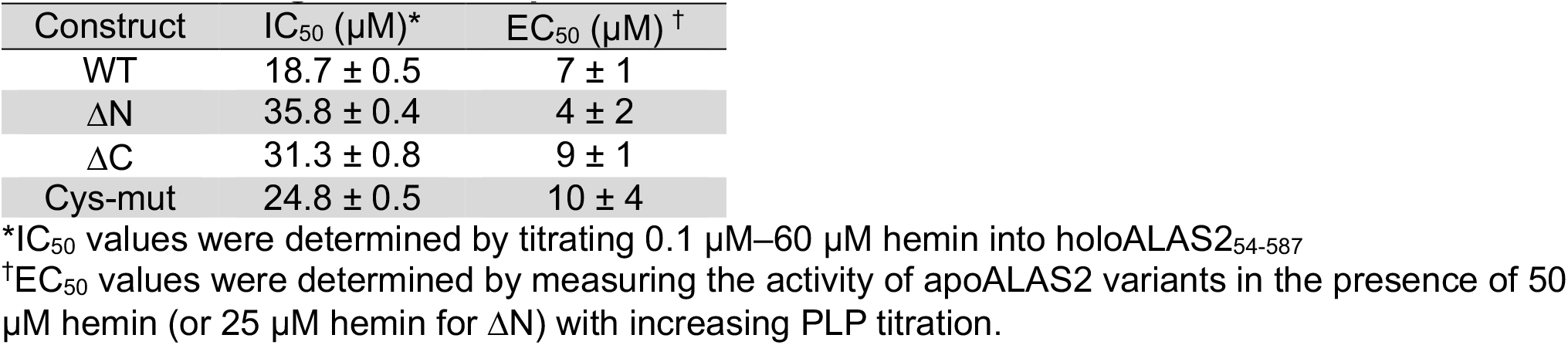
ALAS2-ligand dose-response values.

**Figure 2.**
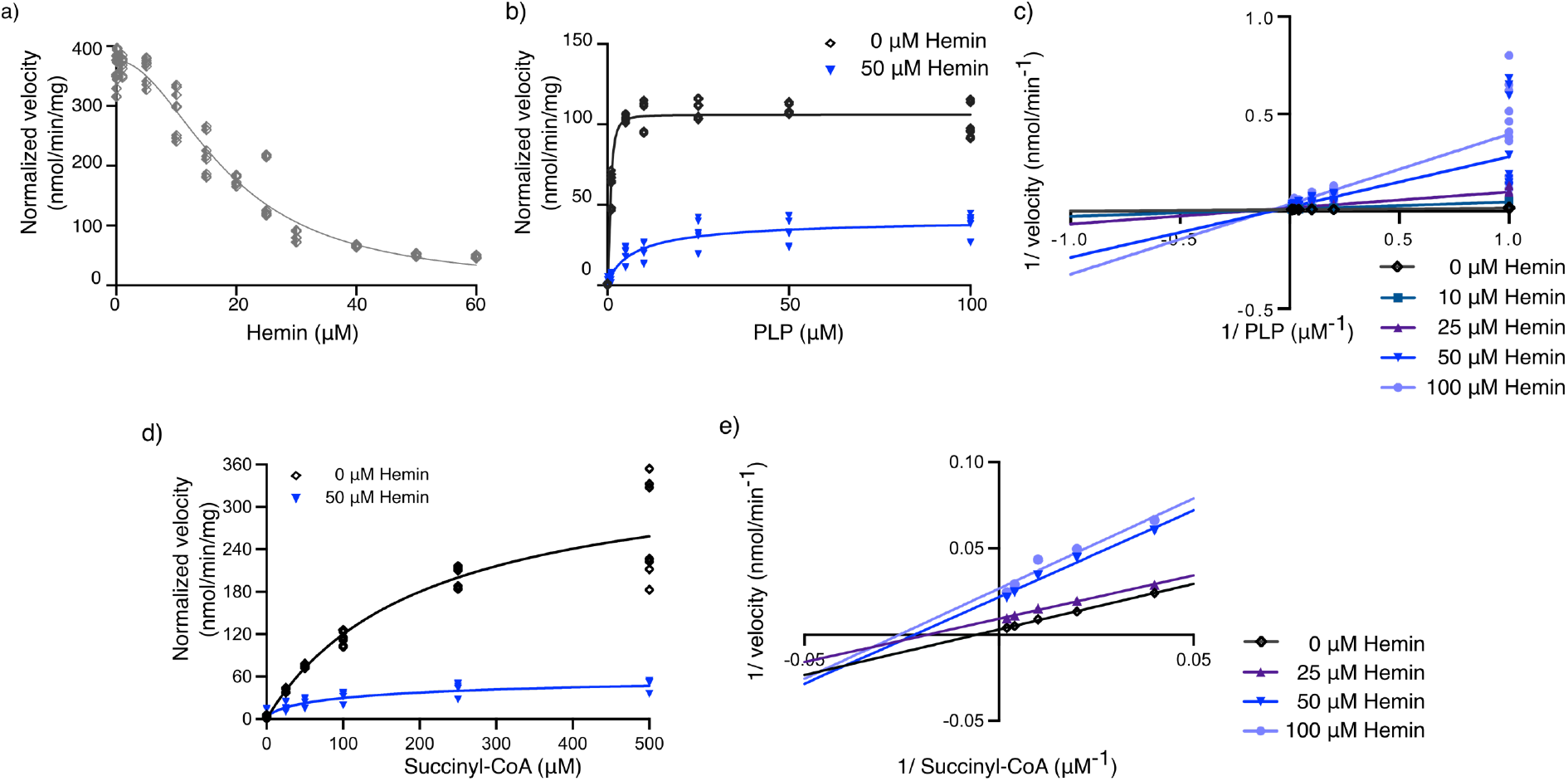
Heme reversibly inhibits the enzymatic activity of human ALAS2. (a) The rate of ALA product released by WT ALAS2 measured as a function of increasing hemin concentration. (b) The rate of ALA production by WT apoALAS2 with increasing concentrations of the PLP cofactor was measured in either the absence (black diamonds) or the presence of 50 μM hemin (blue triangles). (c) Lineweaver-Burk plot depicting the change in enzyme velocity as a function of varied PLP cofactor in the presence of increasing concentrations of the hemin inhibitor (shades of blue). (d) The rate of ALA production by WT holoALAS2 with increasing concentrations of succinyl-CoA was measured in either the absence (black diamonds) or the presence of 50 μM hemin (blue triangles). (e) Lineweaver-Burk plot depicting the change in enzyme velocity as a function of varied succinyl-CoA in the presence of increasing concentrations of the hemin inhibitor (shades of blue).

### Heme binding favors a catalytically inactive ALAS2 conformation

The crystal structure of human ALAS2 lacks the region containing HRM3 and portions of the C-terminal extension due to conformational flexibility and crystallographic disorder^40^. To determine how heme interacts with ALAS2 to inhibit enzyme activity, we used AlphaFold3 (AF3) to model heme bound to the mature ALAS2 homodimer^41^ (**Fig. 3a-b, Supplemental Fig. 3**). The lowest-energy model resulting from the input of two mature ALAS2 chains and two heme b molecules proposes that one of the heme binding sites involves the coordination of two HRMs in the N-terminal and C-terminal extensions (**Fig. 3b-c**). In the absence of inhibitor, these regions are highly flexible, which allows for extensive rearrangement to coordinate heme simultaneously^42,43^. Although the heme-binding site is predicted with lower confidence, all of the models generated by AF3 depict heme coordinated between HRM3 and HRM6 (**Supplemental Fig. 3**). Notably, the manner of heme binding locks the C-terminal extension in a conformation that occludes the active site (**Fig. 3d**). This represents a similar inactive conformation identified in the crystal structure, indicating this orientation is energetically stable^40^. The confidence of the AlphaFold model is not high enough to indicate which of the three possible cysteine residues serve as the axial ligand to coordinate the heme iron center. Also, adding more than two heme molecules in the prediction led to spurious models, indicating the remaining heme binding sites are buried or require a more extensive conformational rearrangement that cannot be accurately modeled.

**Figure 3.**
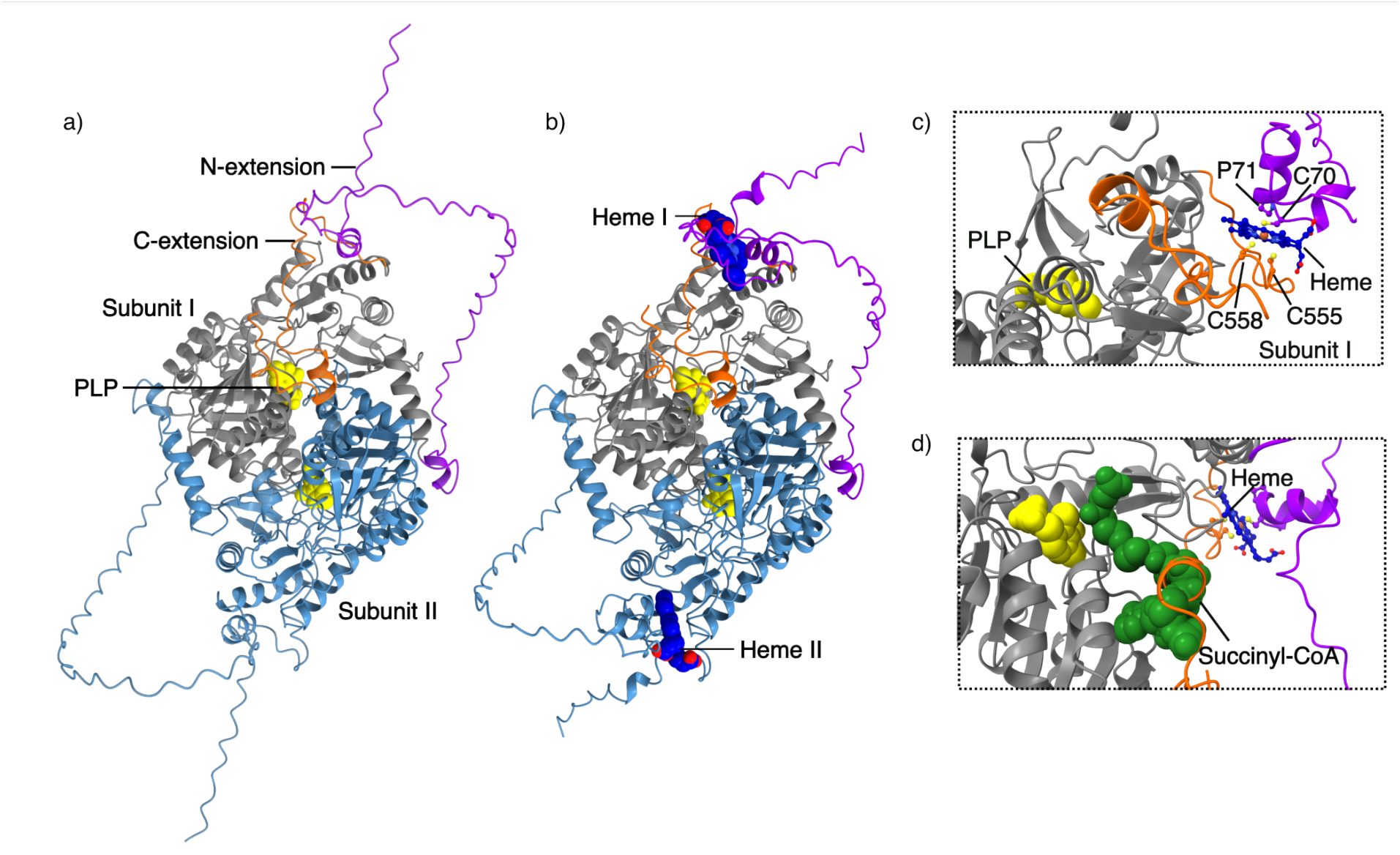
AlphaFold3 model of the ALAS2:heme complex. Two copies of mature ALAS2 (residues 54-587) were modeled without heme (a) and with two heme b molecules (b). The ALAS protomers are colored grey and light blue, the PLP cofactor is shown as yellow spheres, and the heme molecules are shown as blue spheres. The flexible N-terminal extension is colored purple, and the autoinhibitory C-terminal extension is shown in orange. (c) Magnified view of one heme-binding sites reveals heme coordinated between HRM3 (C70-P71) and the C-terminal HRM6 (residues 555-558) from the same chain. (d) Addition of the succinyl-CoA substrate (green spheres) reveals the heme-bound model adopts an inactive conformation where the C-terminal extension sterically interferes with substrate binding. The PLP cofactor and succinyl-CoA substrate were modeled based on superposition with *Rhodobacter capsulatus* ALAS (PDB 2BWO).

### Heme potentially binds at multiple sites to inhibit ALAS2 activity

Multiple ALAS2 constructs with various HRM mutations were generated to interrogate the accuracy of the AlphaFold model and to identify additional heme regulatory sites. These included a truncation without HRM3 (ΔN, residues 75-587), a truncation without the C-terminal CXXC motif (ΔC, residues 54-547), and a construct where all cysteines in these two motifs were mutated to alanines (**Fig. 4a**, Cys-mut). All four constructs were purified as holoenzymes and confirmed to be catalytically active. To determine whether ALAS2 binds hemin when one or more of the HRMs are compromised, ITC was performed with each variant (**Fig. 4b, Table 1**). Titration of ΔN, ΔC, and Cys-mut into hemin resulted in an exothermic binding curve similar to WT. The WT enzyme exhibited the highest affinity for hemin, while disruption of the HRM at either terminus reduced affinity, with a smaller effect observed in the absence of HRM6. Mutation of HRMs at both termini resulted in the lowest affinity for hemin. The ΔN construct lacking HRM3 exhibited an enthalpically driven binding mechanism and a decrease in disorder leading to less efficient binding. This implies that HRM3 plays a prominent role in heme binding compared to the other HRMs. However, other heme-binding sites appear to exist, as revealed by the residual heme binding to our HRM variant constructs.

**Figure 4.**
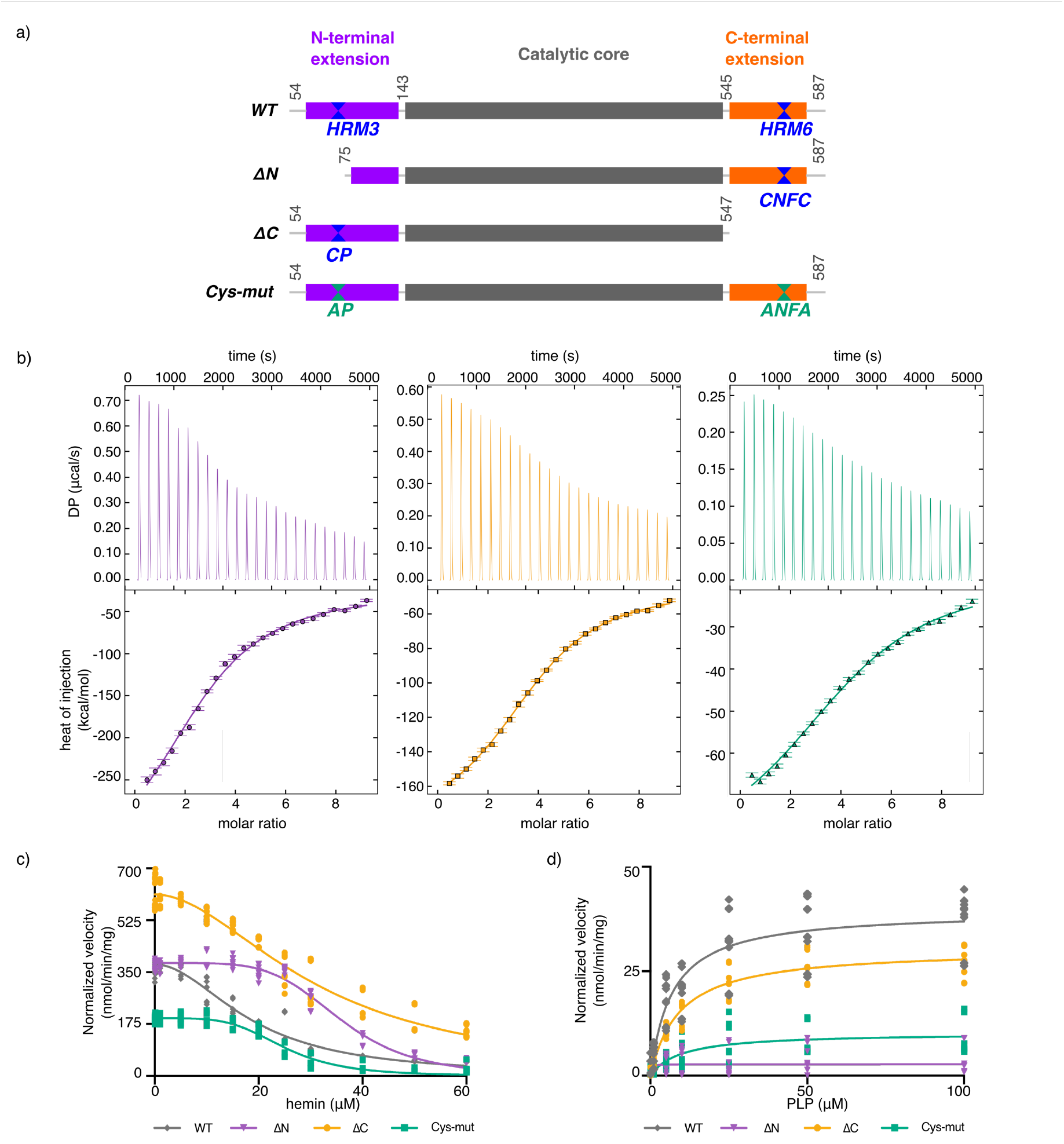
Heme binds human ALAS2 at multiple sites. (a) Domain map of mature ALAS2 WT (colored as in Fig. 1) and HRM variants: WT (residues 54-587), ΔN (residues 75-587), ΔC (residues 54-547), and Cys-mut (C70A/C555A/C558A). (b) Representative ITC thermographs (top) and binding isotherms (bottom) for the interaction between hemin and ALAS2 ΔN (purple, left), ΔC (orange, middle) and Cys-mut (green, right). (c) The rate of ALA release was measured as a function of increasing hemin concentrations for WT (gray diamonds), ΔN (purple triangles), ΔC (orange circles), and Cys-mut (green squares) to monitor inhibition of holoenzyme. (d) The rate of ALA release was measured in the presence of either 50 μM hemin (WT, ΔC, Cys-mut) or 25 μM hemin (ΔN) and increasing concentrations of PLP to determine reactivation of apoenzyme.

Compared to WT ALAS2, the basal activity of the Cys-mut construct decreased by approximately 45%. In contrast, the truncated constructs, ΔN and ΔC, showed activity increases of 5% and 70%, respectively (**Fig. 4c**). The large increase in basal activity of ALAS2 ΔC is expected as the deletion of the C-terminal extension is known to underlie the gain-of-function disorder XLPP^18^. Despite the changes in maximal enzyme velocity, the addition of heme resulted in significant inhibition for all constructs (**Fig. 4c**). Enzymatic activity for all mutant constructs decreased with increasing heme concentrations, but displayed higher IC_50_ values than WT (**Fig. 4c, Table 2**). Among the HRM variants, ALAS2 ΔN was the least sensitive to heme inhibition, exhibiting an IC_50_ value nearly 2-fold higher than WT. Both ΔC and Cys-mut constructs were also inhibited by heme to a lesser extent than WT, with IC_50_ values approximately 1.6 and 1.3 times higher than WT, respectively. There were also notable differences in residual enzyme activity under saturating heme concentrations. The WT, ΔN, and Cys-mut constructs retained approximately 11–13% of basal activity in the presence of 100-fold excess heme. In contrast, ALAS2 ΔC retained ∼25% of basal activity under the same conditions. These results indicate that perturbation of either the N-terminal or C-terminal heme regulatory motifs significantly reduces heme-mediated inhibition, but either motif is sufficient to facilitate ALAS2-heme interaction.

The reversibility of heme inhibition of the HRM variants was monitored by titrating PLP into the apoenzymes preincubated with 50 μM heme (or 25 μM heme for ΔN). In the presence of excess PLP, the apoenzyme activity was restored by ∼50% for WT and ΔC and to over 75% for Cys-mut and ΔN (**Fig. 4d**). Each ALAS2 variant exhibited EC_50_ values comparable to WT (**Table 2**). However, the maximal catalytic activity was restored to varying amounts as both constructs with HRM3 perturbations (ΔN and Cys-mut) displayed the lowest recovery. Additionally, measuring activity in the presence of multiple heme concentrations supported heme interacting as an allosteric mixed inhibitor (**Supplemental Fig. 4**). Overall, these findings provide insight into the reversible interaction between heme and ALAS2, emphasizing its role in regulating enzymatic activity through allosteric effects.

## Discussion

The heme biosynthetic enzyme ALAS2 serves as a critical node, governing the first and rate-limiting step of this essential metabolic pathway. Key differences in ALAS2, including tissue expression, protein interactions, and disease manifestation, necessitate examining this isoform separately from ALAS1.

The interaction between heme and mature human ALAS2 has long been debated but lacked evidence. Here, we provide unprecedented insights into the intricate interaction between heme and the mature human ALAS2 isoform. Our investigation reveals a novel and compelling molecular mechanism of action, wherein heme acts as a potent metabolic inhibitor, effectively decreasing the enzymatic activity of ALAS2. Although labile heme concentrations are difficult to measure directly, estimates vary from less than 1 μM^44^ to greater than 20 μM in humans under non-pathogenic conditions^45^. This upper estimate is commensurate with the IC_50_ values we report for mitochondrial ALAS2-heme inhibition, connecting our *in vitro* research to a plausible cellular state. Critically, this unique mode of inhibition is specific to the enzyme as it localizes to the mitochondrial matrix, where little is known about how ALAS2 responds to chemical cues, but where it primarily operates.

Remarkably, heme binding to ALAS2 exhibits several distinctive traits, including redundancy, reversibility, and allostery. This study supports a model by which multiple motifs act in concert to form a single heme-binding site. Our *in vitro* reconstitution uncovered a process that would be otherwise difficult to identify in cells due to the pleiotropic effects of heme on transcriptional programs and erythropoiesis. The presence of multiple heme-binding sites within the enzyme likely serves as a fail-safe for feedback, ensuring the responsiveness of ALAS2 to heme even if one of the binding sites is compromised. For example, truncations of the ALAS2 C-terminal extension lead to the gain-of-function disease XLPP^18,19,46^, which would affect the enzyme’s ability to respond to heme stress if only one allosteric site existed. However, the presence of multiple non-equivalent heme binding sites across the ALAS2 homodimer makes their isolation and identification difficult, as compensatory changes may occur with enzyme mutagenesis. A redundant method for tuning ALAS2 activity in response to cellular heme levels provides a fitness advantage, and future studies aim to elucidate the unique allosteric properties exhibited by different HRMs.

Structural modeling and analysis reveal that heme functions as an allosteric effector, inducing a conformational change in ALAS2 that prevents substrate binding and, consequently, enzymatic activity. A recent report proposed that heme-bound ALAS2 recruits the CLPXP protease via an adaptor protein to trigger degradation^47^. Similarly, heme binding to the ALAS1 isoform also results in degradation^25,32,33^. Furthermore, our study indicates that heme binding may also influence the interactions of ALAS2 with proteins in the proposed heme synthesis metabolon, a complex of heme biosynthetic enzymes and other mitochondrial proteins that enhance metabolic efficiency^48-50^. This potential modulation of the metabolon would serve as an additional layer of regulation, further influencing the overall efficiency and dynamics of heme production.

Intriguingly, this heme-mediated inhibition discovered in our current work can coexist with a slower, irreversible means of heme-triggered ALAS2 degradation, providing the cell with a multifaceted mechanism to fine-tune heme production in response to various physiological demands (**Fig. 5**).

**Figure 5.**
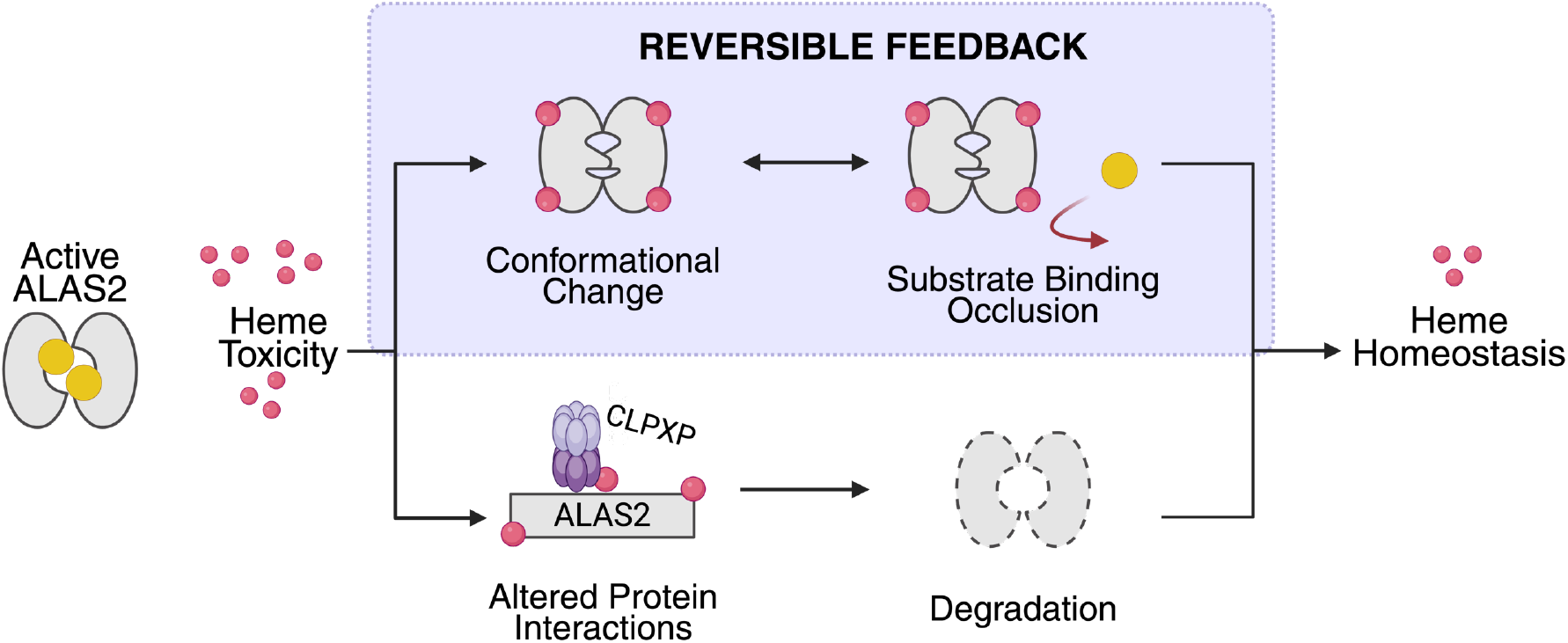
Model of heme-mediated negative feedback inhibition of mitochondrial ALAS2. Under basal conditions, the ALAS2 homodimer (gray ovals) binds PLP (yellow sphere) to subsequently produce heme (red spheres). In the case of toxic heme accumulation, heme binds to ALAS2, with potentially multiple outcomes. We discovered a mechanism where heme binding results in a conformational change that excludes substrate binding (purple box). Additionally, heme binding may recruit mitochondrial proteases, such as CLPXP, for enzyme degradation.

Combined with previous studies of ALAS homologs in bacteria or other tissues^35,51-53^, our work reveals a conserved role in heme-mediated inhibition that has evolved to meet organism and tissue-specific needs. The tight control of heme biosynthesis is paramount, as heme plays a central role in a wide range of cellular processes, including oxygen transport, electron transport, and various metabolic systems such as iron homeostasis and lipid metabolis^54-60^. Given the pivotal role of heme in multiple processes, its dysregulation would also impact physiological functions dependent on other cofactors like FAD and cobalamin, exacerbating disease phenotypes such as anemia, neurotoxicity, oxidative stress, and metabolic dysfunction^63-65^. Therefore, by supporting hemoprotein maturation and integrating with key cellular pathways, the intricate regulation of heme production is crucial for maintaining cellular homeostasis in response to environmental and metabolic demands.

## Experimental Methods

### Protein expression and purification

Human ALAS2 54-587 constructs were cloned into a modified pET28b vector with a ULP1 protease-cleavable N-terminal hexahistidine-SUMO-tag (His_6_-SUMO). The plasmids were transformed into *Escherichia coli* BL21-Codon Plus (DE3)-RIL cells (Agilent technologies) and grown in LB medium containing 25 μg/mL chloramphenicol and 50 μg/mL kanamycin at 37°C and 250 rpm. Expression was induced at an OD_600_ of 0.6-0.8 with 0.5 mM IPTG for 4 hours at 22°C and 250 rpm. Cultures were harvested by centrifugation and the pelleted cells were stored at −80°C.

Cells were resuspended in lysis buffer containing 25 mM HEPES pH 8.0, 400 mM NaCl, 100 mM KCl, 2 mM MgCl_2_, 20 mM imidazole, 10% glycerol, 1 mM DTT, and 20 μM PLP with EDTA-free protease inhibitor (Thermo Fisher Scientific). Cells were lysed by high-pressure homogenization and the lysate was cleared by centrifugation (at 4°C for 30 minutes, 30,000*g*). The clear lysate was incubated with Ni^2+^-NTA agarose resin (Qiagen), pre-equilibrated in lysis buffer, for 1 hour at 4°C. After washing the resin with lysis buffer, the protein was eluted with lysis buffer containing 250 mM imidazole. The His-SUMO tag was cleaved using ULP1 overnight at 4°C while dialyzing into a buffer containing 25 mM HEPES pH 7.0, 100 mM KCl, 10% glycerol, 1 mM DTT, and 20 μM PLP. Tag-free protein was then passed through Ni^2+^-NTA resin pre-equilibrated with dialysis buffer to remove any remaining tagged protein. The protein was further purified with gel-filtration chromatography (HiLoad 16/600 Superdex 200 pg) equilibrated with 25 mM HEPES pH 7, 150 mM KCl, 10% glycerol, and 0.5 mM tris(2-carboxyethyl)phosphine. The purified protein was concentrated using a centrifugal filter (Amicon) flash-frozen in liquid nitrogen, and stored at −80°C.

### Site-directed Mutagenesis

Site-directed mutagenesis was performed on the mature ALAS2 (residues 54–587) construct to generate truncations and cysteine mutants targeting HRM3 and HRM6 (**Supplemental Table 1**). All constructs were verified with Sanger sequencing and were expressed and purified as described above.

### Multiple Sequence Alignment

The amino acid sequences of human (Homo sapiens; P13196, P22557), bovine (Bos taurus; A6QLI6, Q3ZC31) rat (Rattus norvegicus; P13195, Q63147), mouse (Mus musculus; Q8VC19, P08680), and chicken (Gallus gallus: P07997, P18080) ALAS1 and 2 were retrieved from the UniProt database. The multiple sequence alignment was performed using Clustal Omega 1.2.4 with default parameters to identify conserved regions among the sequences. Heme regulatory motif 3 and 6 along with residues flanking the motifs were subsequently used to generate a sequence frequency plot using WebLogo 3.7.4. The alignment of all sequences was visualized using ESPript 3.0.24.

### Isothermal Titration Calorimetry

Binding curves of the hemin-protein interaction were determined using NanoAffinity (TA instruments) at 25°C. Hemin was dissolved in 1 M NaOH and autoclaved to prepare a stock solution of 746 μM, ε_385nm_ = 58,440 M^-1^ cm^-1^. The stock solution was diluted to 1.5 μM in titration buffer containing 25 mM HEPES pH 7, 150 mM KCl, 0.5 mM TCEP. Protein samples were dialyzed into the titration buffer and diluted to a final concentration of 40 μM. Fixed volumes of protein were injected at 200-second intervals into the cell containing hemin. An A+B hetero-association model was used to determine the thermodynamic parameters for each variant. Data were integrated and analyzed using NITPIC and SEDPHAT, and visualized using GUSSI^61-63^.

### Enzyme activity assay

A discontinuous colorimetric activity assay adapted from Shoolingin-Jordan et al. and Bailey et al. was used to measure the enzyme activity^40,64^. The reaction was initiated by adding 100 nM purified enzyme to 50 mM potassium phosphate pH 7.0, 10 mM MgCl_2_, 1 mM DTT, 100 mM glycine and 300 μM succinyl-CoA (175 μL total). The reaction was incubated at 37°C for 15 minutes and terminated with 100 μL of cold 10% trichloroacetic acid. The precipitated protein was removed by centrifugation for 5 minutes, 13,000*g* and 240 μL of the supernatant was mixed with 240 μL of 1 M sodium acetate pH 4.6. To derivatize ALA, 20 μL of acetylacetone was added to the mixture, and boiled at 100°C for 10 minutes. After the sample was cooled for 20 minutes, 100 μL aliquots were transferred into a clear 96-well plate in triplicate. An equal volume of Ehrlich’s reagent (p-Dimethylaminobenzaldehyde, DMAB) was added to each well, and the absorbance was monitored at 553 nm using a microplate reader (Thermo Fisher Scientific). The peak absorbance value was converted to molar quantities of ALA with ε_553_=of 60,400 M^-1^cm^-1^.

Inhibition of enzyme activity was determined by titrating hemin (0.1 μM–60 μM) into the substrate mixture before initiating the reaction upon enzyme addition. Hemin was dissolved in 100% DMSO, ε_403nm_=170,000 M^-1^ cm^-1^, and serially diluted in 100% DMSO to prepare stock solutions for each concentration. The final DMSO concentration in the reaction mixture was 2%. All experiments were performed with a minimum of three biological replicates, each with three technical replicates. Data were fitted using non-linear regression to generate an inhibitor dose-response curve (IC_50_) or an agonist dose-response curve (EC_50_), and statistical significance was determined using a one-way ANOVA with GraphPad Prism 10.

For inhibitor competition experiments, the apoenzymes were preincubated with hemin or PLP (0-100 μM) for 15 minutes at 25°C in the dark before initiating the reaction. Next, either PLP or hemin (0-100 μM) was titrated into the reactions to measure enzyme activation after hemin inhibition or enzyme inhibition of the PLP-treated apoenzyme, respectively. For succinyl-CoA:hemin competition analyses, the WT holoenzyme was mixed with hemin (0, 25, 50, and 100 μM), followed by the addition of increasing succinyl-CoA concentrations (0, 25, 50, 100, 250, and 500 μM). All experiments were performed with a minimum of three biological replicates, each with three technical replicates. Data were fit using non-linear regression to generate an agonist dose-response curve (EC_50_) or an inhibitor dose-response curve (IC_50_). Kinetic parameters (V_max_ and K_M_) were determined using a Michaelis-Menten model in GraphPad Prism 10 and used for generating Lineweaver-Burk Plots.

### In silico modeling

The mature ALAS2-heme model was generated using AlphaFold3 v3.0.0. Two copies of mature ALAS2 (residues 54-587, Uniprot ID P22557) and two copies of heme b ligand were entered as the search query. Of the five models generated, the predicted structure with the highest scores was chosen to generate the heme-bound model. Structure figures were generated using ChimeraX.

## Supplementary material

The supplemental material contains four figures and one table including entire ALAS enzyme multiple sequence alignment, AlphaFold models and corresponding pLDDT scores, Lineweaver Burk plots for HRM variants, and primer sequences used for site-directed mutagenesis.

## Acknowledgements

This work was supported by the National Institute of General Medical Sciences of the National Institutes of Health grant DP2GM146255 (B.L.B.). We thank Lauren P. Jackson for the use of the NanoAffinity ITC instrument. Succinyl-CoA was synthesized by the Vanderbilt Institute for Chemical Biology Chemical Synthesis Core. Biophysical data collection and analysis were provided through the use of the Vanderbilt University Center for Structural Biology Labs and Instrumentation Facility (CSB-LIF), which is an institutionally supported core. Certain figures were created in BioRender.

## Conflict of interest statement

The authors declare no conflict of interest.

## Author Contributions

Iva Chitrakar: Conceptualization, methodology, investigation, validation, formal analysis, visualization, writing

Alexis B. Roberson: Investigation Pedro H. Ayres Galhardo: Investigation

Breann L. Brown: Conceptualization, methodology, investigation, validation, formal analysis, visualization, writing, funding acquisition

## Notes

### Competing Interest Statement

The authors have declared no competing interest.

### Summary of Updates

Current manuscript version includes updated Figure 2.

## References

1. Baker, S.A. & Rutter, J. Metabolites as signalling molecules. Nat Rev Mol Cell Biol 24, 355–374 (2023).

2. Dioum, E.M. et al. NPAS2: a gas-responsive transcription factor. Science 298, 2385–7 (2002).

3. Freeman, S.L. et al. Heme binding to human CLOCK affects interactions with the E-box. Proc Natl Acad Sci U S A 116, 19911–19916 (2019).

4. Yin, L. et al. Rev-erbalpha, a heme sensor that coordinates metabolic and circadian pathways. Science 318, 1786–9 (2007).

5. Dailey, H.A. & Meissner, P.N. Erythroid heme biosynthesis and its disorders. Cold Spring Harb Perspect Med 3, a011676 (2013).

6. Hamza, I. & Dailey, H.A. One ring to rule them all: trafficking of heme and heme synthesis intermediates in the metazoans. Biochim Biophys Acta 1823, 1617–32 (2012).

7. Bechara, E.J.H., Ramos, L.D. & Stevani, C.V. 5-Aminolevulinic acid: A matter of life and caveats. Journal of Photochemistry and Photobiology 7, 100036 (2021).

8. Chiabrando, D., Vinchi, F., Fiorito, V., Mercurio, S. & Tolosano, E. Heme in pathophysiology: a matter of scavenging, metabolism and trafficking across cell membranes. Front Pharmacol 5, 61 (2014).

9. Hunter, G.A., Zhang, J. & Ferreira, G.C. Transient kinetic studies support refinements to the chemical and kinetic mechanisms of aminolevulinate synthase. J Biol Chem 282, 23025–35 (2007).

10. Fratz, E.J. et al. Human Erythroid 5-Aminolevulinate Synthase Mutations Associated with X-Linked Protoporphyria Disrupt the Conformational Equilibrium and Enhance Product Release. Biochemistry 54, 5617–31 (2015).

11. Gibson, K.D., Laver, W.G. & Neuberger, A. Initial stages in the biosynthesis of porphyrins. 2. The formation of delta-aminolaevulic acid from glycine and succinyl-coenzyme A by particles from chicken erythrocytes. Biochem J 70, 71–81 (1958).

12. Kikuchi, G., Kumar, A., Talmage, P. & Shemin, D. The enzymatic synthesis of delta-aminolevulinic acid. J Biol Chem 233, 1214–9 (1958).

13. Bishop, D.F. Two different genes encode delta-aminolevulinate synthase in humans: nucleotide sequences of cDNAs for the housekeeping and erythroid genes. Nucleic Acids Res 18, 7187–8 (1990).

14. Bishop, D.F., Henderson, A.S. & Astrin, K.H. Human delta-aminolevulinate synthase: assignment of the housekeeping gene to 3p21 and the erythroid-specific gene to the X chromosome. Genomics 7, 207–14 (1990).

15. Riddle, R.D., Yamamoto, M. & Engel, J.D. Expression of delta-aminolevulinate synthase in avian cells: separate genes encode erythroid-specific and nonspecific isozymes. Proc Natl Acad Sci U S A 86, 792–6 (1989).

16. Besur, S., Hou, W., Schmeltzer, P. & Bonkovsky, H.L. Clinically important features of porphyrin and heme metabolism and the porphyrias. Metabolites 4, 977–1006 (2014).

17. Stenson, P.D. et al. The Human Gene Mutation Database (HGMD((R))): optimizing its use in a clinical diagnostic or research setting. Hum Genet 139, 1197–1207 (2020).

18. Whatley, S.D. et al. C-terminal deletions in the ALAS2 gene lead to gain of function and cause X-linked dominant protoporphyria without anemia or iron overload. Am J Hum Genet 83, 408–14 (2008).

19. Balwani, M. Erythropoietic Protoporphyria and X-Linked Protoporphyria: pathophysiology, genetics, clinical manifestations, and management. Mol Genet Metab 128, 298–303 (2019).

20. Lee, S., Lee, S., Desnick, R., Yasuda, M. & Lai, E.C. Noncanonical role of ALAS1 as a heme-independent inhibitor of small RNA-mediated silencing. Science 386, 1427–1434 (2024).

21. Lathrop, J.T. & Timko, M.P. Regulation by heme of mitochondrial protein transport through a conserved amino acid motif. Science 259, 522–5 (1993).

22. Munakata, H. et al. Role of the heme regulatory motif in the heme-mediated inhibition of mitochondrial import of 5-aminolevulinate synthase. J Biochem 136, 233–8 (2004).

23. Srivastava, G. et al. Regulation of 5-aminolevulinate synthase mRNA in different rat tissues. J Biol Chem 263, 5202–9 (1988).

24. Peoc’h, K. et al. Regulation and tissue-specific expression of delta-aminolevulinic acid synthases in non-syndromic sideroblastic anemias and porphyrias. Mol Genet Metab 128, 190–197 (2019).

25. Nomura, K. et al. Heme-dependent recognition of 5-aminolevulinate synthase by the human mitochondrial molecular chaperone ClpX. FEBS Lett 595, 3019–3029 (2021).

26. Tanimura, N. et al. Mechanism governing heme synthesis reveals a GATA factor/heme circuit that controls differentiation. EMBO Rep 17, 249–65 (2016).

27. Cox, T.C., Bawden, M.J., Martin, A. & May, B.K. Human erythroid 5-aminolevulinate synthase: promoter analysis and identification of an iron-responsive element in the mRNA. EMBO J 10, 1891–902 (1991).

28. Liu, J. et al. Long non-coding RNA-dependent mechanism to regulate heme biosynthesis and erythrocyte development. Nat Commun 9, 4386 (2018).

29. Campagna, D.R. et al. X-linked sideroblastic anemia due to ALAS2 intron 1 enhancer element GATA-binding site mutations. Am J Hematol 89, 315–9 (2014).

30. Kaneko, K. et al. Identification of a novel erythroid-specific enhancer for the ALAS2 gene and its loss-of-function mutation which is associated with congenital sideroblastic anemia. Haematologica 99, 252–61 (2014).

31. Surinya, K.H., Cox, T.C. & May, B.K. Identification and characterization of a conserved erythroid-specific enhancer located in intron 8 of the human 5-aminolevulinate synthase 2 gene. J Biol Chem 273, 16798–809 (1998).

32. Tian, Q. et al. Lon peptidase 1 (LONP1)-dependent breakdown of mitochondrial 5-aminolevulinic acid synthase protein by heme in human liver cells. J Biol Chem 286, 26424–30 (2011).

33. Kubota, Y. et al. Novel Mechanisms for Heme-dependent Degradation of ALAS1 Protein as a Component of Negative Feedback Regulation of Heme Biosynthesis. J Biol Chem 291, 20516–29 (2016).

34. Zhang, L. et al. Cloning of two 5-aminolevulinic acid synthase isozymes HemA and HemO from Rhodopseudomonas palustris with favorable characteristics for 5-aminolevulinic acid production. Biotechnol Lett 35, 763–8 (2013).

35. Ikushiro, H. et al. Heme-dependent Inactivation of 5-Aminolevulinate Synthase from Caulobacter crescentus. Sci Rep 8, 14228 (2018).

36. Li, T., Bonkovsky, H.L. & Guo, J.T. Structural analysis of heme proteins: implications for design and prediction. BMC Struct Biol 11, 13 (2011).

37. Humayun, F. et al. A Computational Approach for Mapping Heme Biology in the Context of Hemolytic Disorders. Front Bioeng Biotechnol 8, 74 (2020).

38. Schubert, E. et al. Spectroscopic studies on peptides and proteins with cysteine-containing heme regulatory motifs (HRM). J Inorg Biochem 148, 49–56 (2015).

39. Rathod, D.C., Vaidya, S.M., Hopp, M.T., Kuhl, T. & Imhof, D. Shapes and Patterns of Heme-Binding Motifs in Mammalian Heme-Binding Proteins. Biomolecules 13(2023).

40. Bailey, H.J. et al. Human aminolevulinate synthase structure reveals a eukaryotic-specific autoinhibitory loop regulating substrate binding and product release. Nat Commun 11, 2813 (2020).

41. Abramson, J. et al. Accurate structure prediction of biomolecular interactions with AlphaFold 3. Nature 630, 493–500 (2024).

42. Na, I., DeForte, S., Stojanovski, B.M., Ferreira, G.C. & Uversky, V.N. Molecular dynamics analysis of the structural and dynamic properties of the functionally enhanced hepta-variant of mouse 5-aminolevulinate synthase. J Biomol Struct Dyn 36, 152–165 (2018).

43. Stojanovski, B.M., Hunter, G.A., Jahn, M., Jahn, D. & Ferreira, G.C. Unstable reaction intermediates and hysteresis during the catalytic cycle of 5-aminolevulinate synthase: implications from using pseudo and alternate substrates and a promiscuous enzyme variant. J Biol Chem 289, 22915–22925 (2014).

44. Sassa, S. Why heme needs to be degraded to iron, biliverdin IXalpha, and carbon monoxide? Antioxid Redox Signal 6, 819–24 (2004).

45. Aich, A., Freundlich, M. & Vekilov, P.G. The free heme concentration in healthy human erythrocytes. Blood Cells Mol Dis 55, 402–9 (2015).

46. Kadirvel, S. et al. The carboxyl-terminal region of erythroid-specific 5-aminolevulinate synthase acts as an intrinsic modifier for its catalytic activity and protein stability. Exp Hematol 40, 477–86 e1 (2012).

47. Cottle, T., Joh, L., Posner, C., DeCosta, A. & Kardon, J.R. An adaptor for feedback regulation of heme biosynthesis by the mitochondrial protease CLPXP. bioRxiv (2024).

48. Medlock, A.E. et al. Identification of the Mitochondrial Heme Metabolism Complex. PLoS One 10, e0135896 (2015).

49. Grandchamp, B., Phung, N. & Nordmann, Y. The mitochondrial localization of coproporphyrinogen III oxidase. Biochem J 176, 97–102 (1978).

50. Ferreira, G.C., Andrew, T.L., Karr, S.W. & Dailey, H.A. Organization of the terminal two enzymes of the heme biosynthetic pathway. Orientation of protoporphyrinogen oxidase and evidence for a membrane complex. J Biol Chem 263, 3835–9 (1988).

51. Tan, Z. et al. Enhancing thermostability and removing hemin inhibition of Rhodopseudomonas palustris 5-aminolevulinic acid synthase by computer-aided rational design. Biotechnol Lett 41, 181–191 (2019).

52. He, G. et al. Construction of 5-aminolevulinic acid synthase variants by cysteine-targeted mutation to release heme inhibition. J Biosci Bioeng 134, 416–423 (2022).

53. Scholnick, P.L., Hammaker, L.E. & Marver, H.S. Soluble-aminolevulinic acid synthetase of rat liver. II. Studies related to the mechanism of enzyme action and hemin inhibition. J Biol Chem 247, 4132–7 (1972).

54. Dohi, Y. et al. Bach1 inhibits oxidative stress-induced cellular senescence by impeding p53 function on chromatin. Nat Struct Mol Biol 15, 1246–54 (2008).

55. Igarashi, K. & Sun, J. The heme-Bach1 pathway in the regulation of oxidative stress response and erythroid differentiation. Antioxid Redox Signal 8, 107–18 (2006).

56. Zenke-Kawasaki, Y. et al. Heme induces ubiquitination and degradation of the transcription factor Bach1. Mol Cell Biol 27, 6962–71 (2007).

57. Keel, S.B. et al. A heme export protein is required for red blood cell differentiation and iron homeostasis. Science 319, 825–8 (2008).

58. Miller, M.A. et al. CO recombination in cytochrome c peroxidase: effect of the local heme environment on CO binding explored through site-directed mutagenesis. Biochemistry 29, 1777–91 (1990).

59. Vinchi, F. et al. Heme exporter FLVCR1a regulates heme synthesis and degradation and controls activity of cytochromes P450. Gastroenterology 146, 1325–38 (2014).

60. Zhuang, J. et al. Evaluating the roles of the heme a side chains in cytochrome c oxidase using designed heme proteins. Biochemistry 45, 12530–8 (2006).

61. Keller, S. et al. High-precision isothermal titration calorimetry with automated peak-shape analysis. Anal Chem 84, 5066–73 (2012).

62. Zhao, H., Piszczek, G. & Schuck, P. SEDPHAT--a platform for global ITC analysis and global multi-method analysis of molecular interactions. Methods 76, 137–148 (2015).

63. Brautigam, C.A. Calculations and Publication-Quality Illustrations for Analytical Ultracentrifugation Data. Methods Enzymol 562, 109–33 (2015).

64. Shoolingin-Jordan, P.M., LeLean, J.E. & Lloyd, A.J. Continuous coupled assay for 5-aminolevulinate synthase. Methods Enzymol 281, 309–16 (1997).

