## Supplemental Information for "A reversible feedback mechanism regulating mitochondrial heme synthesis"

Supplementary material includes two figures and one table.

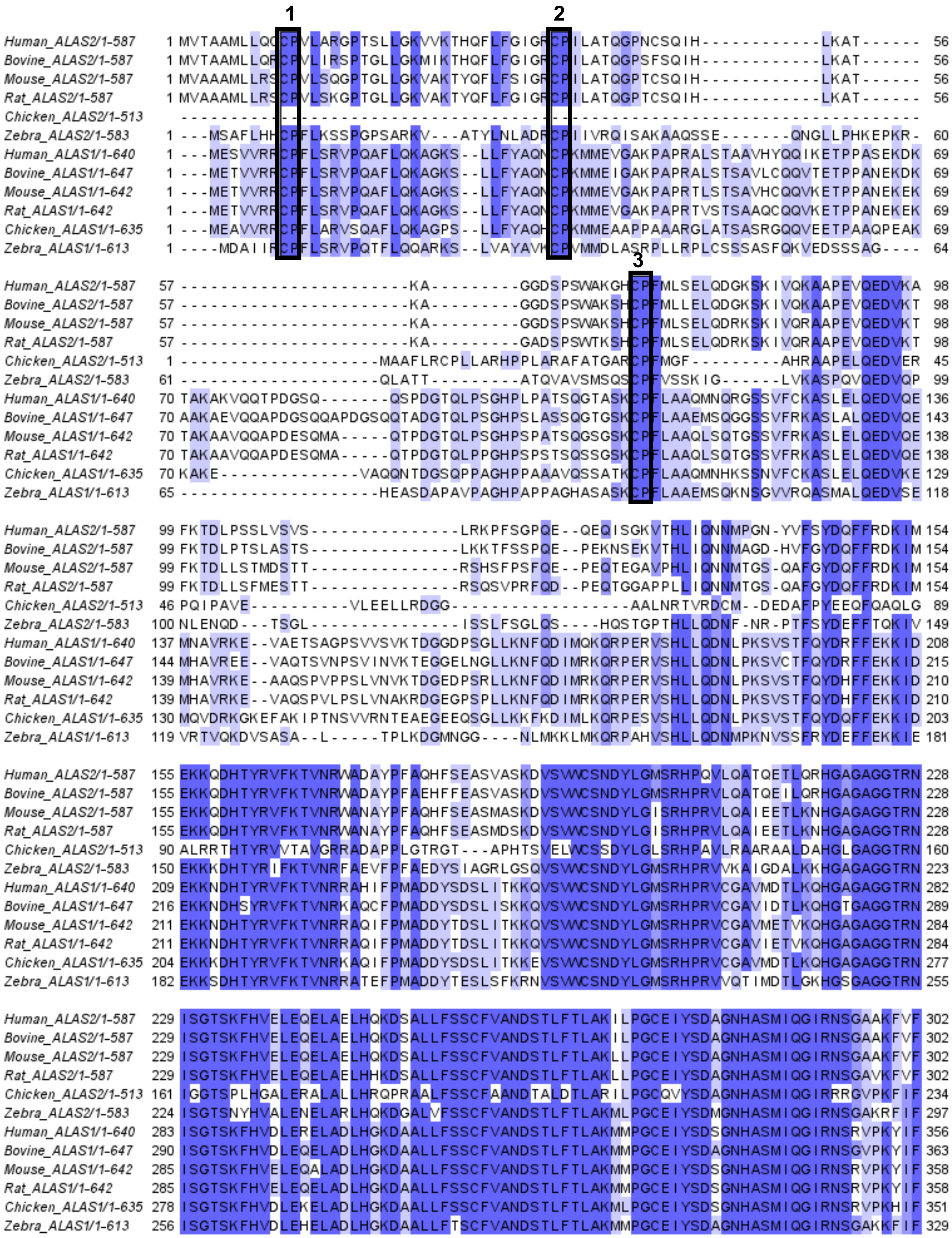
1
2
3
Human_ALAS2/1-587 1 MVTAAMLLQCCPVLARGPTSLLGKVVKTHQFLFGIGRCPILATQGPNCSQIH------------LKAT------ 56
Bovine_ALAS2/1-587 1 MVTAAMLLQRCPVLIRSPTGLLGKMIKTHQFLFGIGRCPILATQGPSFSQIH------------LKAT------ 56
Mouse_ALAS2/1-587 1 MVAAAMLLRSCPVLSQGPTGLLGKVAKTYQFLFSIGRCPILATQGPTCSQIH------------LKAT------ 56
Rat_ALAS2/1-587 1 MVAAAMLLRSCPVLSKGPTGLLGKVAKTYQFLFGIGRCPILATQGPTCSQIH------------LKAT------ 56
Chicken_ALAS2/1-513 --------------------------------------------------------------------------
Zebra_ALAS2/1-583 1 ---MSAFLHHCPFLKSSPGPSARKV---ATYLNLADRCPIIVRQISAKAAQSSE-------QNGLLPHKEPKR- 60
Human_ALAS1/1-640 1 ---MESVVRRCPFLSRVPQAFLQKAGKS--LLFYAQNCPKMMEVGAKPAPRALSTAAVHYQQIKETPPASEKDK 69
Bovine_ALAS1/1-647 1 ---METVVRRCPFLSRVPQAFLQKAGKS--LLFYAQNCPKMMEIGAKPAPRALSTSAVLCQQVTETPPANEKDK 69
Mouse_ALAS1/1-642 1 ---METVVRRCPFLSRVPQAFLQKAGKS--LLFYAQNCPKMMEVGAKPAPRTLSTSAVHCQQVKETPPANEKEK 69
Rat_ALAS1/1-642 1 ---METVVRRCPFLSRVPQAFLQKAGKS--LLFYAQNCPKMMEVGAKPAPRTVSTSAAQCQQVKETPPANEKEK 69
Chicken_ALAS1/1-635 1 ---MEAVVRRCPFLARVSQAFLQKAGPS--LLFYAQHCPKMMEAAPPAAARGLATSASRGQQVEETPAAQPEAK 69
Zebra_ALAS1/1-613 1 ----MDAIIRCPFLSRVPQTFLQQARKS--LVAYAVKCPVMMDLASRPLLRPLCSSSASFQKVEDSSSAG---- 64
Human_ALAS2/1-587 57 -----------------------KA---------GGDSPSWAKGHCPFMLSELQDGKSKIVQKAAPEVQEDVKA 98
Bovine_ALAS2/1-587 57 -----------------------KA---------GGDSPSWAKSHCPFMLLELQDGKSKIVQKAAPEVQEDVKT 98
Mouse_ALAS2/1-587 57 -----------------------KA---------GGDSPSWAKSHCPFMLSELQDRKSKIVQRAAPEVQEDVKT 98
Rat_ALAS2/1-587 57 -----------------------KA---------GADSPSWTKSHCPFMLSELQDRKSKIVQRAAPEVQEDVKT 98
Chicken_ALAS2/1-513 1 --------------------MAAFLRCPLLARHPPLARAFATGARCPFMGF---------AHRAAPELQEDVER 45
Zebra_ALAS2/1-583 61 --------------------QLATT----------ATQVAVSMSQSCPFVSSKIG------LVKASPQVQEDVQP 99
Human_ALAS1/1-640 70 TAKAKVQQTPDGSQ-------QSPDGTQLPSGHPLPATSQGTASKCPFLAAQMNQRGSSVFCKASLELQEDVQE 136
Bovine_ALAS1/1-647 70 AAKAEVQQAPDGSQQAPDGSQQTADGTQLPSGHPSLASSQGTGSKCPFLAAEMSQGGSSVFRKASLALQEDVQE 143
Mouse_ALAS1/1-642 70 TAKAAVQQAPDESQMA-----QTPDGTQLPSGHPSPATSQGSGSKCPFLAAQLSQTGSSVFRKASLELQEDVQE 138
Rat_ALAS1/1-642 70 TAKAAVQQAPDESQMA-----QTPDGTQLPPGHPSPSTSQSSGSKCPFLAAQLSQTGSSVFRKASLELQEDVQE 138
Chicken_ALAS1/1-635 70 KAKE--------------VAQQNTDGSQPPAGHPPAAAVQSSATKCPFLAAQMNHKSSNVFCKASLELQEDVKE 129
Zebra_ALAS1/1-613 65 --------------------HEASDAPAVPAGHPAPPAGHASASKCPFLAAEMSQKNSGVVRQASMALQEDVSE 118
Human_ALAS2/1-587 99 FKTDLPSSLVSVS---------------LRKPFSGPQE--QEQISGKVTHLIQNNMPGN-YVFSYDQFFRDKIM 154
Bovine_ALAS2/1-587 99 FKTDLPTSLASTS---------------LKKTFSSPQE--PEKNSEKVTHLIQNNMAGD-HVFGYDQFFRDKIM 154
Mouse_ALAS2/1-587 99 FKTDLLSTMDSTT---------------RSHSFPSFQE--PEQTEGAVPHLIQNNMTGS-QAFGYDQFFRDKIM 154
Rat_ALAS2/1-587 99 FKTDLLSFMESTT---------------RSQSVPRFQD--PEQTGGAPPLLIQNNMTGS-QAFGYDQFFRDKIM 154
Chicken_ALAS2/1-513 46 PQIPAVE----------VLEELLRDGG-----------------AALNRTVRDCM--DEDAFPYEEQFQAQLG 89
Zebra_ALAS2/1-583 100 NLENQD---TSGL---------------ISSLFSGLQS----HQSTGPTHLLQDNF-NR-PTFSYDEFFTQKIV 149
Human_ALAS1/1-640 137 MNAVRKE--VAETSAGPSVVSVKTDGGDPSGLLKNFQDIMQKQRPERVSHLLQDNLPKSVSTFQYDRFFEKKID 208
Bovine_ALAS1/1-647 144 MHAVREE--VAQTSVNPSVINVKTEGGELNGLLKNFQDIMRKQRPERVSHLLQDNLPKSVCTFQYDRFFEKKID 215
Mouse_ALAS1/1-642 139 MHAVRKE--AAQSPVPPSLVNVKTDGEDPSRLLKNFQDIMRKQRPERVSHLLQDNLPKSVSTFQYDHFFEKKID 210
Rat_ALAS1/1-642 139 MHAVRKE--VAQSPVLPSLVNAKRDGEGPSPLLKNFQDIMRKQRPERVSHLLQDNLPKSVSTFQYDHFFEKKID 210
Chicken_ALAS1/1-635 130 MQVDRKGKEFAKIPTNSVVRNTEAEGEEQSGLLKKFKDIMLKQRPESVSHLLQDNLPKSVSTFQYDQFFEKKID 203
Zebra_ALAS1/1-613 119 VRTVQKDVSASA--L-----TPLKDGMNGG----NLMKKLMKQRPAHVSHLLQDNMPKNVSSFRYDEFFEKKIE 181
Human_ALAS2/1-587 155 EKKQDHTYRVFKTVNRWADAYPFAQHFSEASVASKDVSVWCSNDYLGMSRHPQVLQATQETLQRHGAGAGGTRN 228
Bovine_ALAS2/1-587 155 EKKQDHTYRVFKTVNRWADAYPFAEHFFEASVASKDVSVWCSNDYLGMSRHPRVLQATQEILQRHGAGAGGTRN 228
Mouse_ALAS2/1-587 155 EKKQDHTYRVFKTVNRWANAYPFAQHFSEASMASKDVSVWCSNDYLGISRHPRVLQAIEETLKNHGAGAGGTRN 228
Rat_ALAS2/1-587 155 EKKQDHTYRVFKTVNRWANAYPFAQHFSEASMDSKDVSVWCSNDYLGISRHPRVLQAIEETLKNHGAGAGGTRN 228
Chicken_ALAS2/1-513 90 ALRRTHTYRVVTAVGRRADAPPLGTRGT---APHTSVELWCSSDYLGLSRHPAVLRAARAALDAHGLGAGGTRN 160
Zebra_ALAS2/1-583 150 EKKKDHTYRIFKTVNRFAEVFPFAEDYSIAGRLGSQVSVWCSNDYLGMSRHPRVVKAIGDALKKHGAGAGGTRN 223
Human_ALAS1/1-640 209 EKKNDHTYRVFKTVNRRAHIFPMADDYSDSLITKKQVSVWCSNDYLGMSRHPRVCGAVMDTLKQHGAGAGGTRN 282
Bovine_ALAS1/1-647 216 EKKNDHSYRVFKTVNRKAQCFPMADDYSDSLISKKQVSVWCSNDYLGMSRHPRVCGAVIDTLKQHGTGAGGTRN 289
Mouse_ALAS1/1-642 211 EKKNDHTYRVFKTVNRRAQIFPMADDYTDSLITKKQVSVWCSNDYLGMSRHPRVCGAVMETVKQHGAGAGGTRN 284
Rat_ALAS1/1-642 211 EKKNDHTYRVFKTVNRRAQIFPMADDYTDSLITKKQVSVWCSNDYLGMSRHPRVCGAVIETVKQHGAGAGGTRN 284
Chicken_ALAS1/1-635 204 EKKKDHTYRVFKTVNRKAQIFPMADDYSDSLITKKEVSVWCSNDYLGMSRHPRVCGAVMDTLKQHGAGAGGTRN 277
Zebra_ALAS1/1-613 182 EKKSDHTYRVFKTVNRRATEFPMADDYTESLSFKRNVSVWCSNDYLGMSRHPRVVQTIMDTLGKHGSGAGGTRN 255
Human_ALAS2/1-587 229 ISGTSKFHVELEQELAELHQKDSALLFSSCFVANDSTLFTLAKILPGCEIYSDAGNHASMIQGIRNSGAAKFVF 302
Bovine_ALAS2/1-587 229 ISGTSKFHVELEQELAELHQKDSALLFSSCFVANDSTLFTLAKILPGCEIYSDAGNHASMIQGIRNSGAAKFVF 302
Mouse_ALAS2/1-587 229 ISGTSKFHVELEQELAELHQKDSALLFSSCFVANDSTLFTLAKLLPGCEIYSDAGNHASMIQGIRNSGAAKFVF 302
Rat_ALAS2/1-587 229 ISGTSKFHVELEQELAELHHKDSALLFSSCFVANDSTLFTLAKLLPGCEIYSDAGNHASMIQGIRNSGAVKFVF 302
Chicken_ALAS2/1-513 161 IGGTSPLHGALERALALLHRQPRAALFSSCFAANDTALDTLARILPGCQVYSDAGNHASMIQGIRRRGVPKFIF 234
Zebra_ALAS2/1-583 224 ISGTSNYHVALENELARLHQKDGALVFSSCFVANDSTLFTLAKMLPGCEIYSDMGNHASMIQGIRNSGAKRFIF 297
Human_ALAS1/1-640 283 ISGTSKFHVDLERELADLHGKDAALLFSSCFVANDSTLFTLAKMMPGCEIYSDSGNHASMIQGIRNSRVPKYIF 356
Bovine_ALAS1/1-647 290 ISGTSKFHVDLEQELADLHGKDAALLFSSCFVANDSTLFTLAKMMPGCEIYSDAGNHASMIQGIRNSGVPKYIF 363
Mouse_ALAS1/1-642 285 ISGTSKFHVELEQALADLHGKDAALLFSSCFVANDSTLFTLAKMMPGCEIYSDSGNHASMIQGIRNSRVPKYIF 358
Rat_ALAS1/1-642 285 ISGTSKFHVELEQELADLHGKDAALLFSSCFVANDSTLFTLAKMMPGCEIYSDSGNHASMIQGIRNSRVPKYIF 358
Chicken_ALAS1/1-635 278 ISGTSKFHVDLEKELADLHGKDAALLFSSCFVANDSTLFTLAKMLPGCEIYSDSGNHASMIQGIRNSRVPKHIF 351
Zebra_ALAS1/1-613 256 ISGTSKFHVDLEHELADLHGKDAALLFTSCFVANDSTLFTLAKMMPGCEIYSDAGNHASMIQGIRNSGAKKFIF 329

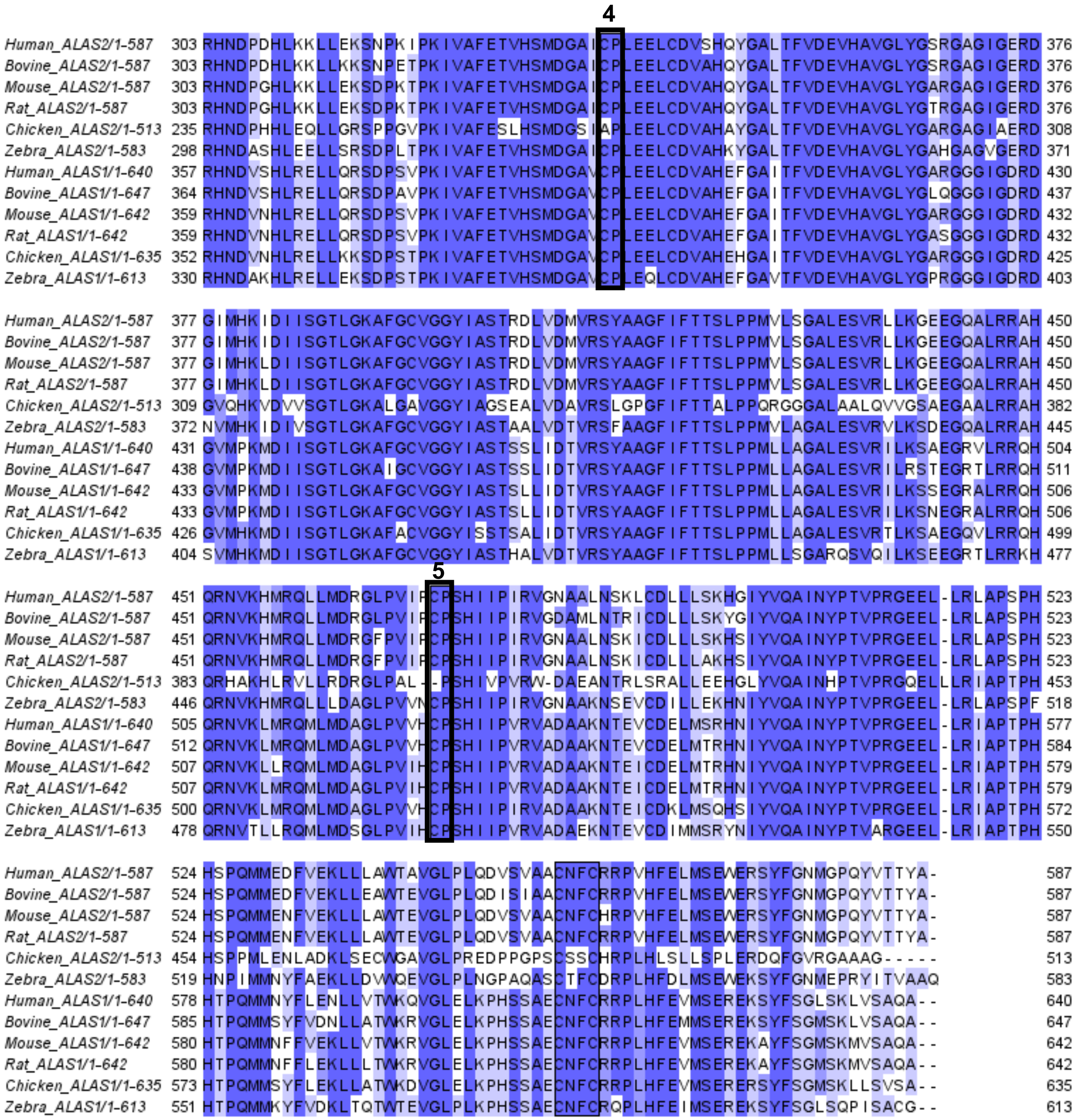


**Supplemental Figure 1.** Sequence alignment of ALAS homologs from various vertebrate organisms. Identical residues are shaded in dark blue, similar residues are shaded in light blue, and the heme regulatory motifs (HRMs) are highlighted in black boxes and numbered from N- to C-terminus.

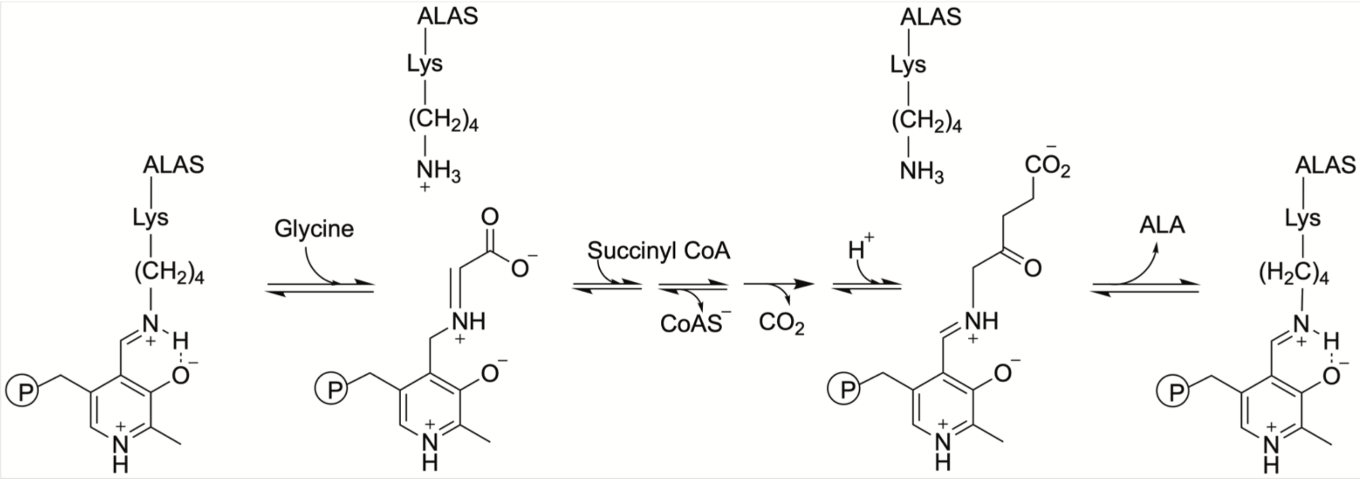


PLP – Lys PLP – Glycine PLP – ALA PLP – Lys

**Supplemental Figure 2.** PLP-dependent catalytic mechanism of ALAS. PLP initially forms an internal aldimine by covalently binding a conserved lysine residue in ALAS (PLP-Lys, Lys 391 for human ALAS2). Upon glycine binding, the covalent pyridoxal lysine bond is broken, and the PLP is converted to an external aldimine (PLP-Glycine). Reaction with succinyl-CoA yields an intermediate that undergoes decarboxylation and condensation to form 5-aminolevulinic acid (ALA), which is released to regenerate the internal aldimine.

a)

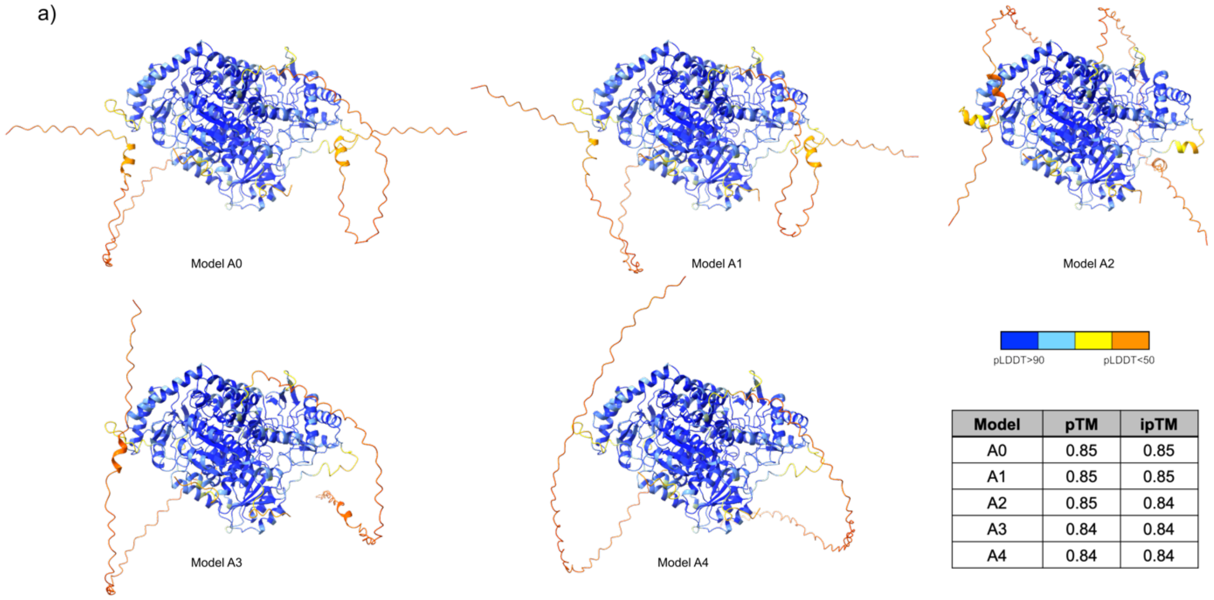


| Model | pTM | ipTM |
|---|---|---|
| A0 | 0.85 | 0.85 |
| A1 | 0.85 | 0.85 |
| A2 | 0.85 | 0.84 |
| A3 | 0.84 | 0.84 |
| A4 | 0.84 | 0.84 |

b)

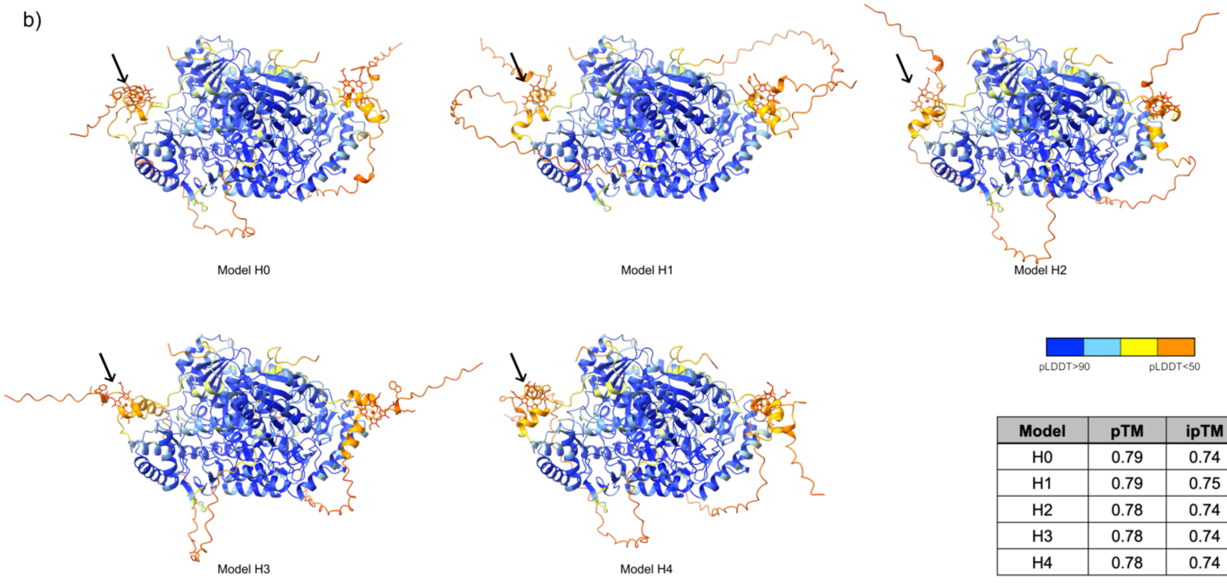


| Model | pTM | ipTM |
|---|---|---|
| H0 | 0.79 | 0.74 |
| H1 | 0.79 | 0.75 |
| H2 | 0.78 | 0.74 |
| H3 | 0.78 | 0.74 |
| H4 | 0.78 | 0.74 |

**Supplemental Figure 3.** AlphaFold3 models and associated confidence scores. The top five models output from AlphaFold3 modeling of (a) two mature human ALAS2 chains (residues 54-587) alone, and (b) two mature hALAS2 chains with two heme b molecules (stick representation, black arrows). The models are colored according to pLDDT score, where higher values represent higher model confidence. In all models, the heme molecule is docked between HRM3 ($^{70}$CP$^{71}$) and HRM6 ($^{555}$CNFC$^{558}$). The predicted template modeling (pTM) score and the interface predicted template modeling (ipTM) score are shown in the inset table.

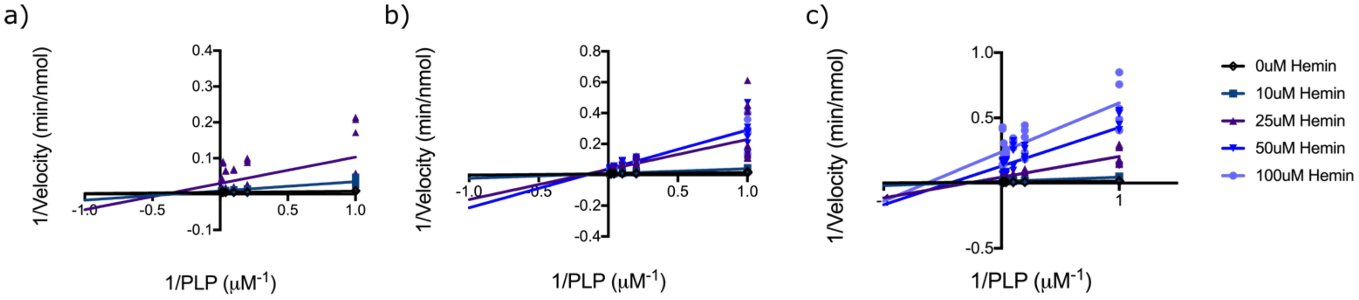


**Supplemental Figure 4.** Lineweaver Burk plots of ALAS2 a) ΔN, b) ΔC and c) Cys-mut depicting the change in reciprocal enzyme velocity with increasing concentrations of the hemin inhibitor (shades of blue).

**Supplemental Table 1. Primer sequences for generating human ALAS2 constructs**

| Construct | Primer sequence (5'→3') |
|---|---|
| hALAS2$_{54\text{-}587}$ (WT) | Forward: CAGACCGGTGGTAAGGCAACAAAGGCTGGA<br>Reverse: GTCACCACCTATGCCTGAGCGGCCGCAGGACC |
| hALAS2$_{75\text{-}587}$ (ΔN) | Forward: AGGACCACCGGTGGCTCGGAACTCCAGGATGG<br>Reverse: GTCACCACCTATGCCTGAGCGGCCGCAGGACC |
| hALAS2$_{54\text{-}547}$ (ΔC) | Forward: CAGACCGGTGGTAAGGCAACAAAGGCTGGA<br>Reverse: GGTGGGGCTGCCCCTCTAAGCGGCCGCAGGACC |
| hALAS2$_{54\text{-}587}$ C(Cys-mut) | Forward (C70A): GCGAAGGGCCACGCTCCCTTCATGCTG<br>Reverse (C70A): CAGCATGAAGGGAGCGTGGCCCTTCGC<br>Forward (C555A/C558A): TGTGGCTGCCGCGAATTTCGCGCGCCGTCCTG<br>Reverse (C555A/C558A): GCAGCCACAGACACATCCTGG |
